# Preclinical evaluation of a natural extract-based oral nanoformulation from *Eucalyptus tereticornis* for potential use in treating type 2 diabetes mellitus

**DOI:** 10.64898/2026.04.16.719114

**Authors:** Natalia Arbeláez, Elkin Escobar-Chaves, Andrea Correa, Adriana Restrepo, Sergio Acin, Jahir Orozco, Norman Balcazar

**Affiliations:** PECET Group, Faculty of Medicine. Universidad de Antioquia. Medellín, Colombia; GENMOL Group, Faculty of Natural and Exact Sciences. Universidad de Antioquia. Medellín, Colombia; Department of Physiology and Biochemistry, Faculty of Medicine. Universidad de Antioquia, Calle 70, N° 52-21, A.A. 1226, Medellín, Colombia; Max Planck Tandem Group in Nanobioengineering, University of Antioquia. Complejo Ruta N, Calle 67 N° 52–20, Medellín, 050010, Colombia

**Keywords:** Toxicity, preclinical study, polymeric nanocarrier, T2DM, triterpenes, *Eucalyptus tereticornis*

## Abstract

The acute, subacute, and subchronic oral toxicities, as well as the combined chronic toxicity and carcinogenicity, of a nanotechnology-based formulation derived from a natural extract of *Eucalyptus tereticornis* leaves were investigated. This nanoformulation demonstrates anti-obesogenic and potentially anti-diabetic properties. Our study aims to conduct preclinical tests to evaluate the chemical formulation. To assess acute toxicity, rats received a single oral dose of 2000 mg/kg of the nanoformulation. In the subacute trial, mice were treated with approximately 1180 mg/kg of the nanoformulation for 28 days. In the combined chronic toxicity and carcinogenicity study, the nanoformulation was administered daily at approximately 590 mg/kg for 10 months. At the end of the experiment, hematological, biochemical, and histopathological assessments were conducted. Throughout the acute, subacute, subchronic, and chronic/carcinogenicity studies, animals showed no toxic effects from the treatment or the vehicle. No histopathological lesions, such as degeneration or cell death in the liver, kidney, or gastrointestinal tract, were observed. Treatments did not cause any clinical changes, and there were no significant differences in weight, hematological, or biochemical parameters. Therefore, the nanoformulation did not produce toxic effects in the animals.

## Introduction

Type 2 diabetes mellitus (T2DM) has reached epidemic proportions. It is estimated that about 589 million adults had diabetes mellitus in 2024, and the prevalence is projected to reach 853 million by 2050. Increased intake of high-calorie diets and reduced physical activity have contributed to this rise and will likely continue to do so (1). Obesity is the most significant metabolic disorder involved in developing insulin resistance and the potential onset of T2DM. Between 60% and 90% of cases of T2DM are attributable to obesity [1,2].

Several treatments exist for T2DM, but limited efficacy, side effects, and the need for intravenous administration drive the global search for new, more effective therapeutic agents. Natural products have long been used to prevent or treat various diseases, and interest in researching, developing, and marketing drugs, nutraceuticals, and dietary supplements is increasing [3].

Previous studies identified a prototype drug for T2DM—a standardized fraction of triterpenes extracted from the leaves of *Eucalyptus tereticornis*—that, when administered intraperitoneally, reverses metabolic abnormalities in an animal model of prediabetes. It reduces fasting hyperglycemia, glucose and insulin intolerance, total cholesterol, Low-Density Lipoprotein (LDL), Very Low-Density Lipoprotein (VLDL) levels, and blood leptin levels, while significantly decreasing the weight of obese animals. Additionally, findings confirm the effect of a mixture of three triterpenes on fatty acid breakdown in the liver and adipocytes. They also demonstrate anti-inflammatory activity in the adipose tissue of obese mice, which likely helps break the cycle between mild chronic inflammation and insulin resistance [4,5].

To enhance the bioavailability of a natural triterpene-rich extract suitable for oral intake, a nanotechnology-based formulation has been produced in a batch. With particle sizes of 165 nm and 213 nm for empty and loaded NPs, respectively, a polydispersity index below 0.2, and an average zeta potential of -35 mV, it demonstrates effective properties for this purpose. The encapsulation efficiency (EE) exceeds 98% for ursolic acid lactone (LAU) and over 85% for the mixture of ursolic acid (UA) and oleanolic acid (OA), the main triterpenes in the extract. Release tests in physiological media show increased release at pH 7, reaching up to 80% for LAU and 50% for UA+OA after 72 hours, compared to pH 5 [6].

This formulation produces effects similar to those seen with peritoneal administration of the triterpene mixture, including reductions in body and liver weight in diet-induced obese mice with insulin resistance. It also improves glucose and insulin tolerance and reduces intracellular accumulation of neutral fats (triacylglycerols, TAG) in liver tissue. Additionally, it lowers serum TAG and LDL levels [6]. Several preclinical studies have examined the encapsulation of pharmaceutically active ingredients (PAIs) into polymeric nanoparticles. Notable examples include cases like ours, where multiple PAIs are encapsulated, creating synergistic effects, coordinated delivery, and enhanced intracellular permeability. These advantages can lead to increased therapeutic effectiveness, fewer side effects, and lower drug resistance [7]. Some studies use poly(lactic-co-glycolic) acid (PLGA) as a key component in the design of polymeric nanoparticles (NPs) [8], and its potential for drug development has been supported by evidence of safety and efficacy in living organisms [9]. PLGA is a widely used polymer for the manufacture of various types of NPs and is an FDA-approved delivery system for human use. It forms the base polymer in the development of the nanoformulation described herein.

Several studies have confirmed the use of PLGA-based NPs for delivering drugs to treat diabetes mellitus [10], central nervous system diseases [11], certain cancers [12], and vascular disorders [13]. Additionally, other polymers have been used to create NPs that, like PLGA, improve drug efficacy and bioavailability. All these preclinical studies support the use of NPs as drug-delivery vehicles due to their proven effectiveness and safety [14, 15].

This study aims to conduct preclinical safety and efficacy tests on a nanoformulation derived from a natural extract with anti-obesogenic and potentially anti-diabetic properties, in accordance with the Organization for Economic Cooperation and Development (OECD) guidelines for the evaluation of chemical substances. The tests included acute oral toxicity [16], repeated-dose oral toxicity over 28 [17] and 90 days [18], and long-term toxicity combined with carcinogenicity [19].

## Materials and methods

### Preparation of plant extract

Preparation of the crude extract (OBE100) and chromatographic separation of triterpenes from *Eucalyptus tereticornis* leaves were conducted according to the previously established protocol [5]. Briefly, 25 kg of dried E*. tereticornis* leaves were used to extract OBE100 via liquid-liquid extraction with a 4:3:1 hexane:methanol: water mixture (v/v). The organic phase was then collected and filtered under vacuum. The precipitate was finally gathered and stored for future use (5). After chromatographic analysis, the main components of the crude extract were ursolic acid (UA), oleanolic acid (OA), and ursolic acid lactone (UAL) (Fig. 1A).

**Fig 1.**
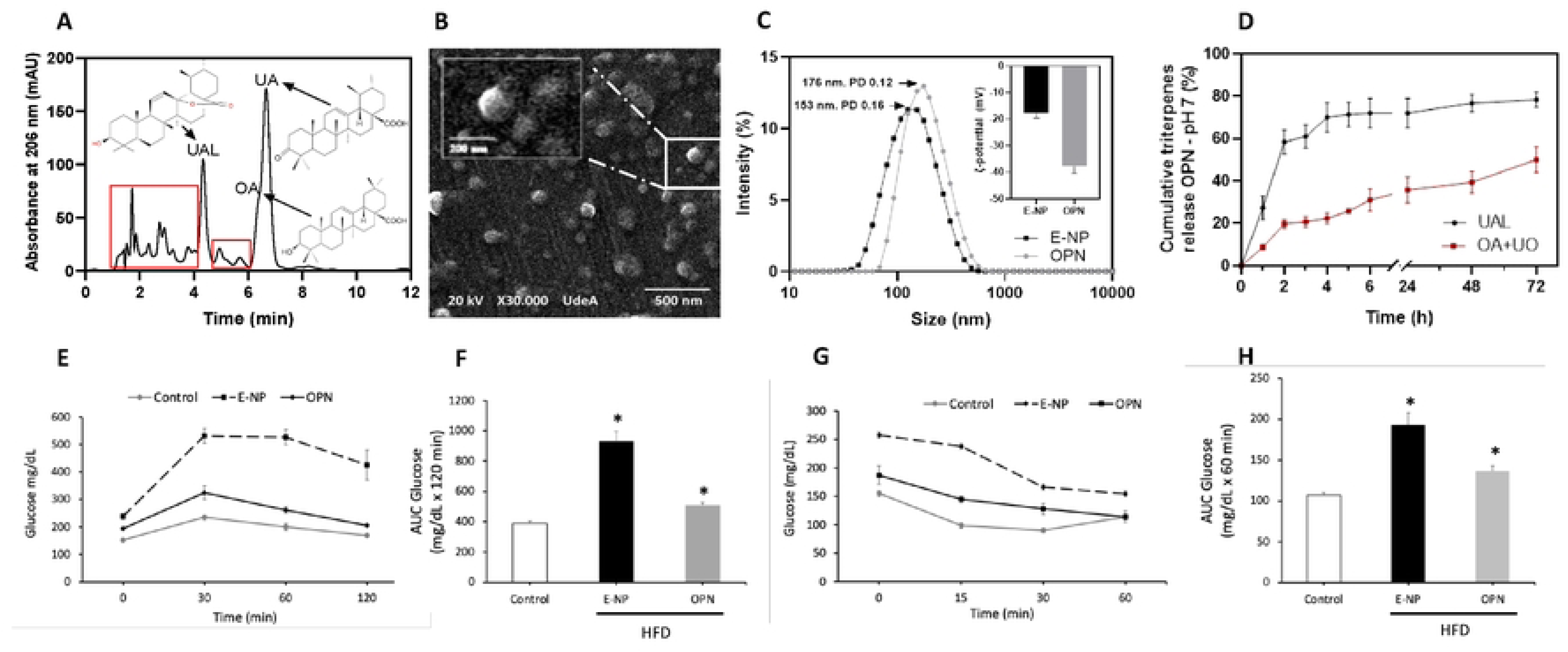
Physicochemical characterization and release profile of the polymeric nanoformulation. (A) Chromatographic profiles of OBE100’s main triterpenes: ursolic acid lactone (UAL), oleanolic acid (OA), ursolic acid (UA), and minor components (red square). (B) Morphological characterization of OPN by SEM. (C) Hydrodynamic size and ζ-potential of OBE100-loaded nanoparticles (ONP) in grey and empty nanoparticles (NP) in black. (D) Release profile of OBE100-loaded NP: main triterpenes UAL and OA+UO in sink conditions at phosphate buffer pH 7, with N=3. The effect of oral particle nanoformulation (OPN) on carbohydrate metabolism is assessed via (E) glucose tolerance test (IPGTT) and (G) insulin tolerance test (ITT) in prediabetic mice treated orally with 214 mg/kg doses of OBE100-loaded NP (OPN) and empty NP (E-NP). (F) and (H) show the AUCs for glucose and insulin, respectively, calculated from data in (E) and (G) on a high-fat diet (HFD). N=6-8 mice per group.

### Scaling up the production of the oral polymeric nanoformulation (OPN)

To carry out the preclinical studies, a larger-scale production system was developed based on the previously standardized oral polymeric nanoformulation (OPN) [6]. The emulsion/solvent evaporation method was employed, with hydrophobic compounds prepared in the organic phase by dissolving 5 g of PLGA 50:50, 5 g of Kolliphor® P188, 1 g of freeze-dried crude extract OBE100, and 504 mg of D-α-tocopherol in 250 ml of ethyl acetate. Subsequently, 500 mL of water was added to the organic phase. The mixture was divided into 135-mL aliquots, vortexed at 3,200 rpm for 20 seconds, and homogenized with seven ultrasonication pulses, 28 seconds each (on 4 seconds, off 10 seconds), using a ½-inch sonic tip at 65% amplitude (Ultrasonicator Q500, Qsonica, Newton, CT, USA) to create the nanoemulsion and ensure OBE100 encapsulation. Finally, the nanoemulsion was placed in a rotary evaporator at 200 mbar, 150 rpm, at 45°C for 15 minutes (Rotavapor® R-300, Buchi, Germany). The resulting OPN was purified by dialysis against 20% ethanol with a dialysis membrane (MWCO 12-14 kDa, Spectrum™) for 24 hours. The final sample was then freeze-dried for physicochemical characterization tests. This protocol was used to prepare different batches.

### Physicochemical characterization of the large-scale produced oral polymeric nanoformulation

The diameter and polydispersity index (PDI) of the nanoparticles were measured using dynamic light scattering (DLS), and the surface charge (ζ-potential) was determined by electrophoretic light scattering (ELS) with a NanoZs instrument (Malvern Instruments, UK) at 25°C after aqueous dilution in triplicate. To confirm the morphology of the OBE100-loaded nanoparticles, scanning electron microscopy (SEM) and transmission electron microscopy (TEM) were used. For SEM, samples were mounted on graphite tape, coated with a thin layer of gold (Au) using a Denton vacuum desk IV, and analyzed in a high-vacuum SEM (JEOL JSM-6490LV, Akishima, Tokyo, Japan). For TEM, 5 µL of OPN aqueous dispersion (1,5 mg/mL) was drop cast to a copper grid with a carbon film, dried at room temperature for 24 hours in a desiccant silica gel chamber, stained with 1% uranyl acetate for 8 minutes, washed with deionized water, and analyzed in the electron microscope (Tecnai F20 Super Twin, FEI). Encapsulation efficiency (%EE), loading capacity (%LC), and release kinetics were determined by high-performance liquid chromatography (HPLC) according to the previously described protocol [6].

### Assessment of the metabolic effects of the oral polymeric nanoformulation in a mouse model of diet-induced obesity

The OPN prepared in a laboratory setting has already been evaluated (6). The Diet-Induced Obesity (DIO) mouse model was established to verify the biological effects of the batch-prepared material using glucose and insulin tolerance tests, fasting glucose, and lipid profiles. Male C57BL/6J mice over 4 weeks old from the SPF Animal Facility at the University of Antioquia Research Headquarters were used. They were housed at 22±2 °C on a 12-hour light/dark cycle and randomly assigned to two groups: the control group (n=7), which received a standard diet (ND) with 14% fat, 54% carbohydrates, and 32% protein, and the high-fat diet (HFD) group (n=14), which was fed a diet consisting of 42% fat, 42% carbohydrates, and 15% protein, for 10 weeks with free access to food and water.

Once it was confirmed that the HFD-fed groups were hyperglycemic and insulin-resistant, they were divided into a control group treated with empty nanoparticles (E-NP) and another group that received 15 doses of OBE 100 encapsulated into nanoparticles (OPN) at 214 mg/kg of body weight, administered orally. After completing the treatment, fasting glucose, the Insulin Tolerance Test (ITT), and the Intraperitoneal Glucose Tolerance Test (IPGTT) were performed [4]. The Institutional Animal Care and Use Committee of the University of Antioquia approved all animal studies (Protocol number 65).

### Animals and Housing

For the acute oral toxicity study, nine female Wistar rats *(Rattus norvegicus)* were used. These rats were Specific Pathogen Free (SPF), exogamous, nulliparous, and 8 to 12 weeks old, with a maximum body weight variation of 20% among individuals. For other studies, Swiss Webster CFW mice (*Mus musculus),* SPF and exogamous, all obtained from Charles River Laboratories, were employed. The rats were housed in polycarbonate cages measuring 45 cm long, 25 cm wide, and 30 cm high, while the mice were kept in polycarbonate cages measuring 20 cm long, 25 cm wide, and 20 cm high. Both cage types contained 5 cm of sterilized pine bedding, water, and LabDiet® 5010 brand food, available ad libitum. The macroenvironmental conditions included a temperature of 21°C ± 2°C, relative humidity of 40-65% (both maintained by a timer), an air exchange rate of 16 to 20 air changes per hour with High Efficiency Particulate Air (HEPA) filters, artificial white light illumination, and a 12/12-hour light/dark cycle regulated by a timer.

### Preclinical Experimental Procedures

All studies involved animals randomly assigned to three groups: OBE100 encapsulated in nanoparticles (OPN), empty nanoparticles (E-NP), and a control group treated with PBS (Control). The number of animals in each study was as follows: acute oral toxicity according to OECD guideline 423 of 2002 (n = 9 rats total) [16], repeated-dose oral toxicity over 28 days (subacute) following OECD Guideline 407 of 2008 (n = 26 mice total) [17], repeated-dose oral toxicity over 90 days (subchronic) according to OECD guideline 408 of 2018 (n = 52 mice total) [18], and a combined chronic toxicity/carcinogenicity study based on OECD guideline 453 of 2018 (n = 120 mice) [19]. All treatments were administered orally using an 18-G stainless steel orogastric tube for rats and a 21-G stainless steel orogastric tube for mice. The Institutional Animal Care and Use Committee of the University of Antioquia approved the pre-clinical studies (Acta number 140, 01-07-2021).

### Clinical Assessment

In all studies, daily clinical monitoring was conducted alongside treatment administration and weekly body weight measurements, including the two weeks (14 days) after treatment concluded. In the acute oral toxicity study, careful monitoring was conducted for the first 24 hours after treatment. In the repeated-dose/90-day oral toxicity study, an ophthalmological assessment of the anterior chamber was performed at both the beginning and end, along with evaluations of responses to stimuli such as proprioception, threat reflex, and eye reflex. Fourteen days after each study ended, animals were euthanized in a CO₂ chamber, then underwent intracardiac puncture to collect postmortem blood samples, and finally received gross necropsy to examine all organ systems.

### Hematological and Blood Chemistry Parameters

Postmortem blood samples were collected for: complete blood counts in subchronic and chronic studies; liver function tests (ALT, AST) in all studies; and blood ions (sodium, chloride, potassium), as well as kidney function tests (creatinine, BUN, urea) in subacute and subchronic studies.

### Histopathology analysis

During the necropsy, organ samples were collected for histopathological analysis in accordance with the study protocol. The combined chronic and carcinogenesis study examined the stomach, small intestine, colon, esophagus, liver, pancreas, spleen, adrenal gland, seminal vesicle, kidneys, ovaries, uterus, testicles, epididymis, prostate, submandibular lymph nodes, trachea, lungs, brain, cerebellum, heart, skin, submandibular salivary gland, and eyeball. A semi-quantitative method was used to evaluate lesion severity, employing the following scale: 0 = no lesion; 1 = mild lesion; 2 = mild to moderate lesion; 3 = moderate lesion; 4 = moderate to severe lesion; and 5 = severe lesion. The severity scores were compared across the test groups. For acute, subacute, and subchronic toxicity studies, the organs specified in the OECD guidelines were analyzed, and only those with observable changes are reported in the results.

### Statistical analysis

Data are presented as means ± SEM. Comparisons between groups were analyzed using a one-way analysis of variance (ANOVA) followed by a Tukey-Kramer multiple comparison test. Student’s t-tests were used for individual pairwise comparisons of least squares means. The trapezoidal rule was applied to determine the area under the curve (AUC). Differences were considered significant at P < 0.05. Analyses were performed using Prism 10 (GraphPad Software Inc., La Jolla, CA, USA).

## RESULTS

### Scaling, physical characterization, and biological evaluation of the polymeric nanoformulation

The standardized scaling method for producing the oral nanoformulation is reproducible for Extract OBE100, remains stable over time without undergoing aggregation or flocculation, and maintains appropriate sizes for oral administration. Nanometric particles measuring 153 nm for empty-NP with a ζ-potential of -17.9 mV, a PDI of 0.16, and 176 nm for OPN with a ζ-potential of -37.8 mV and a PDI of 0.12 were achieved. These physicochemical properties are consistent with those obtained in batch, thereby demonstrating that nanoparticles with the desired specifications can be reliably scaled up (Fig. 1B and 1C, and Fig. S1). The encapsulation efficiency was 90%, and the loading capacity was 6.16%. The release kinetics align with those observed in previous studies of oral nanoparticle production at the laboratory scale: 70% and 35% of ursolic acid lactone and the ursolic-oleanolic mixture, respectively, were released within the first 6 hours, and up to 80% and 50%, respectively, were released after 72 hours (Fig. 1D).

The OPN was tested using a DIO mouse model, and results were measured as AUC from 0 to 120 minutes for the IPGTT test and from 0 to 60 minutes for the ITT test. After three weeks of various treatments (15 doses), mice fed a high-fat diet (HFD) and treated with the oral nanoformulation (OPN) showed improved glucose tolerance compared to HFD-fed mice without treatment (E-NP). Similar results were seen when analyzing the ABC generated by the ITT tests (Fig. 1E, H). Administering 75.4 mg/kg of OBE100 encapsulated into OPN significantly improved glucose and insulin tolerance in prediabetic animals (Fig. 1), confirming the biological effect of the large-scale-prepared nanoformulation.

### Acute oral toxicity study

The nine rats used in the trial were randomly divided into three experimental groups: OPN group: treated with a single oral dose of 2000 mg/kg of the nanoformulation containing OBE100, the E-NP group: treated with a single dose of 2000 mg/kg of empty NPs, and the control group: given a single dose of 1 ml of PBS.

After administering the different treatments, no clinical signs of disease or toxicity were observed during the first 24 hours and the following 14 days of follow-up. Body weight gain remained stable from day 1 to day 14 of the study (Fig. 2A).

**Fig 2.**
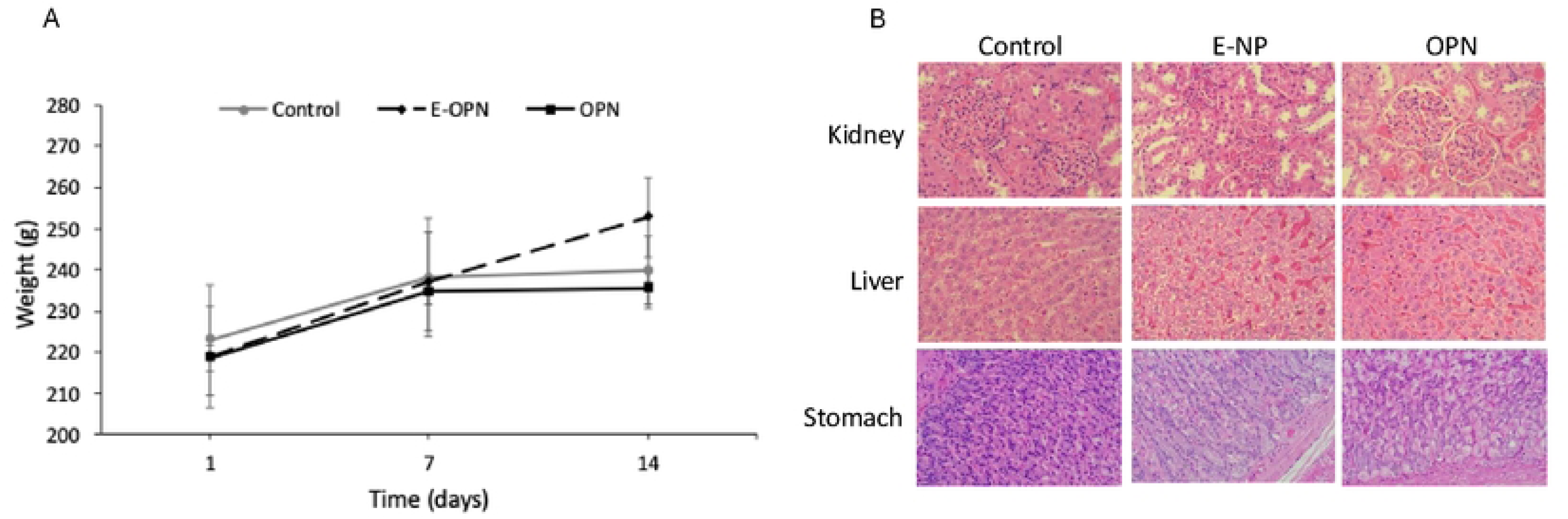
A) Body weight change in the acute oral toxicity study. Time: days after treatment. B) Photomicrographs of the kidneys, liver, and stomach from rats in the three groups of the acute oral toxicity study. The images are at 400X magnification and stained with hematoxylin and eosin (H&E). The control group received PBS; E-NP indicates empty nanoparticles; OPN indicates the oral nanoformulation with OBE100.

During the necropsy, all organ systems were examined, and no gross lesions were found in any organs. Samples from the liver, kidneys, spleen, both glandular and non-glandular stomachs, pancreas, small intestine, and large intestine were collected for histopathological analysis. The liver samples from the three test groups showed hepatocyte karyomegaly and inflammatory foci. These findings do not signify pathological lesions; instead, they are likely due to the strain’s genetic background rather than the study. Membranoproliferative glomerulitis, associated with a genetic background, was observed in the kidney samples. No lesions were identified in the stomach or other organs (Fig. 2B).

### Subacute oral toxicity study

Twenty-six Swiss mice (CFW) were randomly divided into three experimental groups: five females and five males for the OPN and E-NP groups, and three females and three males for the control group with PBS. The OPN group received 300 μl (average 1100 mg/kg) of the OBE100 nanoformulation; the E-NP group was given 300 μl (average 920 mg/kg) of empty NPs. The control group received daily treatment with 300 μl of PBS. All treatments were administered daily for 28 days.

During daily treatment, no clinical abnormalities were observed, and body weight remained stable across all three groups (Table 1 and Fig. 3A). During necropsy, no macroscopic lesions were found in any organs. Samples of the liver, kidney, spleen, glandular and non-glandular stomach, pancreas, and heart were collected for histopathological analysis.

**Fig 3.**
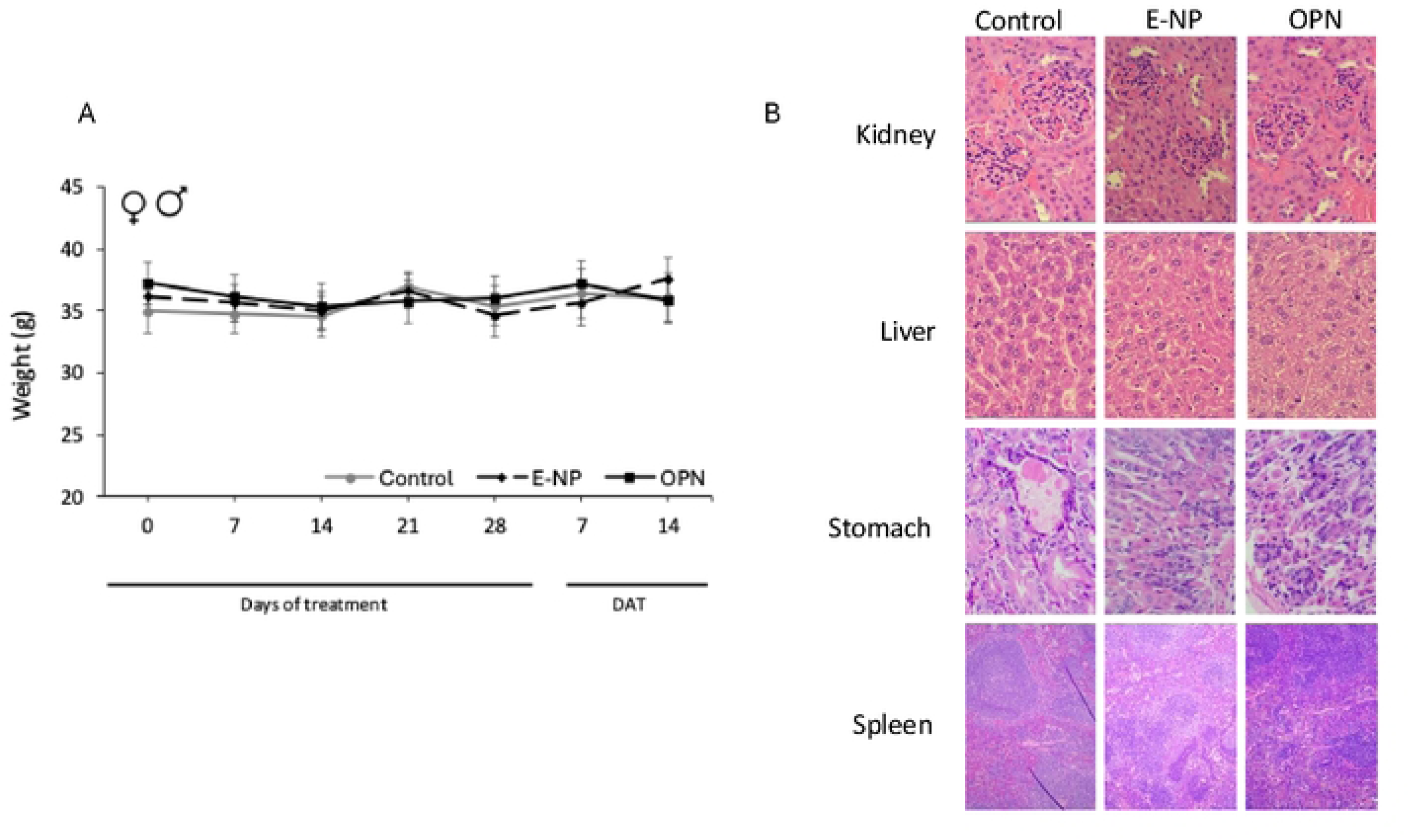
A) Body weight change in the subacute oral toxicity study. DAT: days after treatment. Both sexes are included. B) Photomicrographs of the kidneys, liver, stomach, and spleen from mice in the three groups of the subacute oral toxicity study. The images are at 100X and 400X magnification and stained with hematoxylin and eosin (H&E). The control group received PBS; E-NP denotes empty nanoparticles; OPN denotes the oral nanoformulation with OBE100.

**Table 1.** Blood count and chemistry values in mice across three groups in the subacute oral toxicity study. Control mice received PBS, E-NP (empty nanoparticles), and OPN (oral nanoformulation with OBE100). Data are shown as means ± SD for each group. No statistically significant differences were found in the measured parameters between the experimental groups. Reference values are from references 32 and 36.

|  | Experimental groups |  |  |
| --- | --- | --- | --- |
| Parameters | Control | E-NP | OPN |
| Liver function |  |  |  |
| ALT (40-69 UI/L) | 134,4±52,3 | 81,5±28,1 | 145,8±95,6 |
| AST (86-212 U/L) | 131,2±44,9 | 98,9±22,2 | 342,1±136,4 |
| Plasma Ion Concentration |  |  |  |
| Sodium (112-193 mEq/L) | 154,9±19,6 | 157,5±10,8 | 150,7±9 |
| Chlorine (82-150 mEq/L) | 153,8±24,9 | 129,3±20,4 | 143,3±28,2 |
| Potassium (5,1-10,4 mEq/L) | 13,8±2,2 | 10±0 | 8,8±3,6 |
| Renal Function |  |  |  |
| Creatine (0,3-1 md/dL) | 0,5±0,04 | 0,5±0,02 | 0,6±0,2 |
| BUN (17-28 mg/dL) | 25,3±0 | 22,5±0,4 | 28,2±8,6 |
| Urea (36-67 mg/dL) | 54,3±0 | 48,2±0,9 | 62,8±18,9 |

Liver samples from all three groups showed karyomegaly. They also showed small clusters of inflammatory cells composed of lymphocytes and plasmacytes, as well as extramedullary hematopoiesis in the spleen. These lesions are related to the genetic background and are not considered pathological; they are part of their normal morphology (Fig. 3B). No lesions linked to the toxicity of the administered treatment were observed in histological sections from both the OPN and E-NP groups. Fatty change was slightly more noticeable in the control group animals treated with PBS.

In kidney samples, mild to moderate membranoproliferative and mesangioproliferative glomerulonephritis were observed, with reduced urinary space. This was not related to the treatment but to the strain (Fig. 3B). No tubular changes were evident due to nephrotoxicity or inflammation. The presence of lymphocytes and plasma cells in the interstitium was minimal. No lesions were found in other organs.

Complete blood counts for the three groups were within normal ranges. Blood glucose was measured with the Accu-Chek® glucometer after six hours of fasting, with no abnormalities detected (data not shown). Regarding blood chemistry analysis (Table 1), liver transaminase levels exceeded the upper limits in all three treatment groups (ALT and AST), with slightly higher levels in mice treated with OPN, particularly for AST. However, no micro- or macroscopic lesions attributable to the treatment were observed. No changes were seen in plasma ion concentrations or renal function.

### Sub-chronic oral toxicity study

Fifty-two Swiss mice (CFW) were randomly divided into three experimental groups: ten females and ten males for the OPN and E-NP groups, and six females and six males as controls receiving PBS. The OPN group received 300 μl (average 1100 mg/kg) of the OBE100 nanoformulation; the E-NP group was given 300 μl (average 920 mg/kg) of empty NPs. The control group received daily treatment with 300 μl of PBS. All treatments were administered daily for 90 days.

In the control group, weight remained stable, and greater gains were observed than in the other two groups (Fig. 4A, B). Weight loss in any group did not exceed 10% of the initial weight, so it was not a reason to terminate the experiment for any mouse.

**Fig 4.**
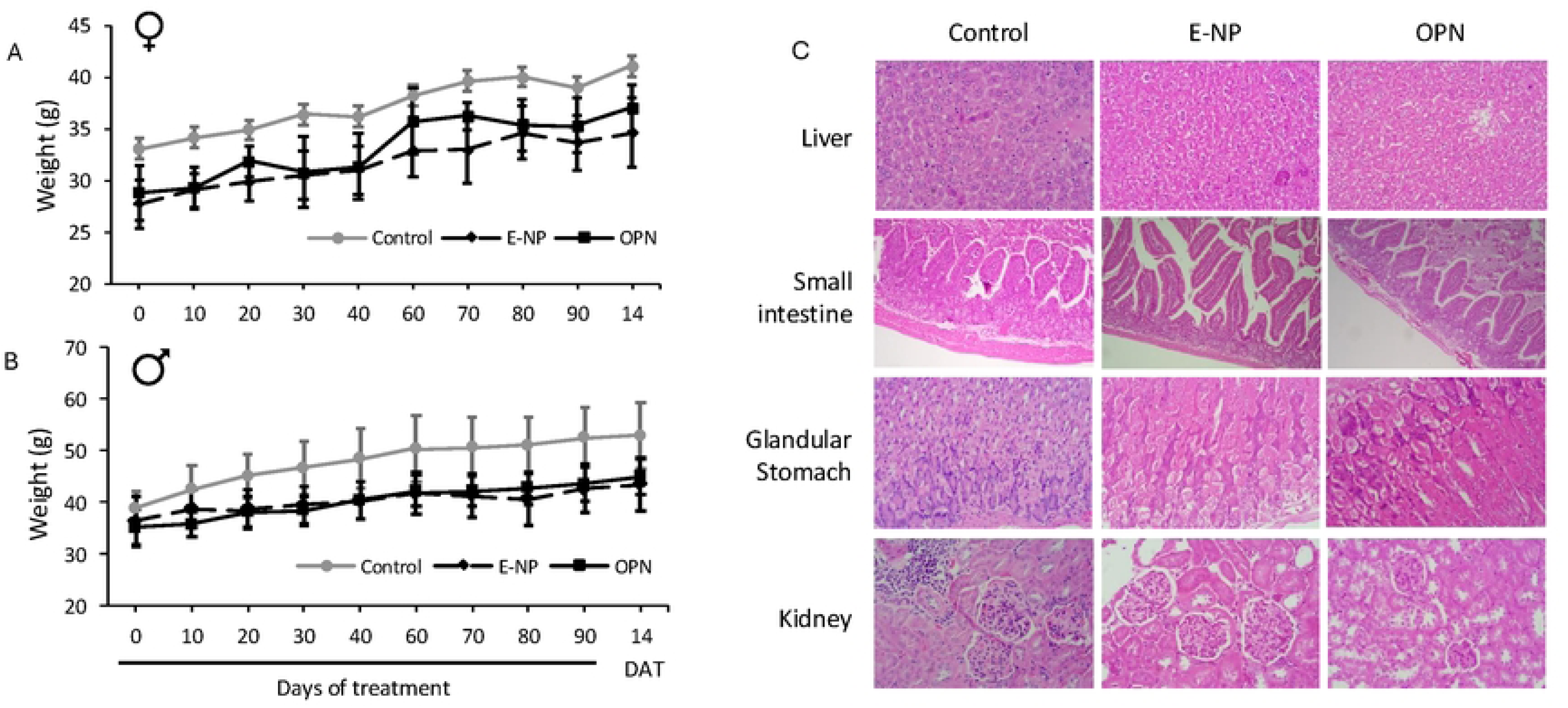
Body weight change during the subchronic oral toxicity study. A. Female, B. Male. DAT: days after treatment. C) Photomicrographs of the kidneys, liver, stomach, and spleen from mice in the three groups of the subchronic oral toxicity study. The images are at 100X and 400X magnification and stained with hematoxylin and eosin (H&E). The control group received PBS; E-NP denotes empty nanoparticles; OPN denotes the oral nanoformulation with OBE100.

Blood counts were within normal ranges for all three study groups. Blood glucose levels after six hours of fasting remained within normal ranges (data not shown), as did plasma ion concentrations and blood chemistry used to assess renal function (Table 2). ALT enzyme levels were slightly elevated in the OPN-treated group, but the difference was not statistically significant; similarly, organ function was not significantly different (Table 2).

**Table 2.**
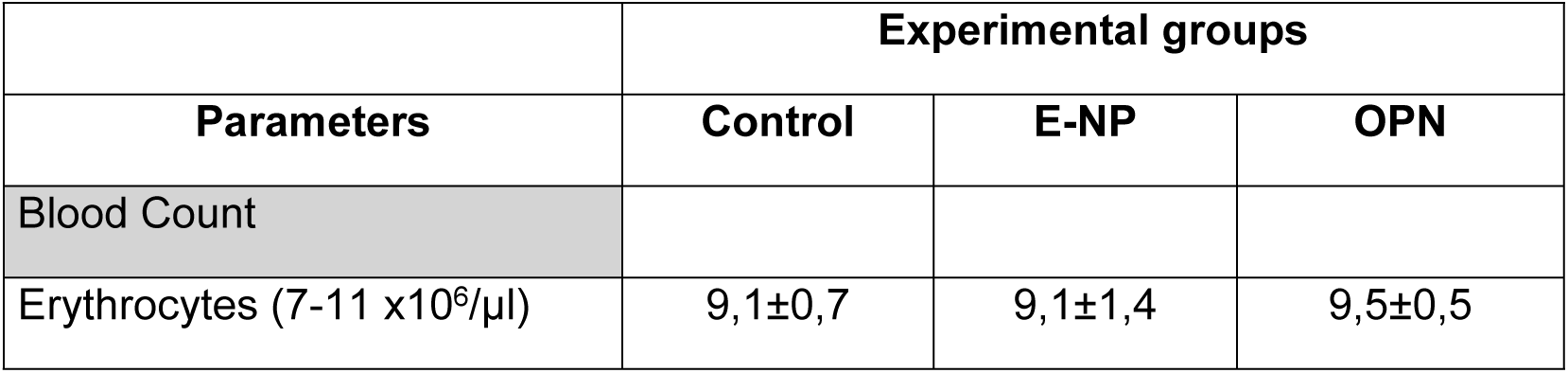
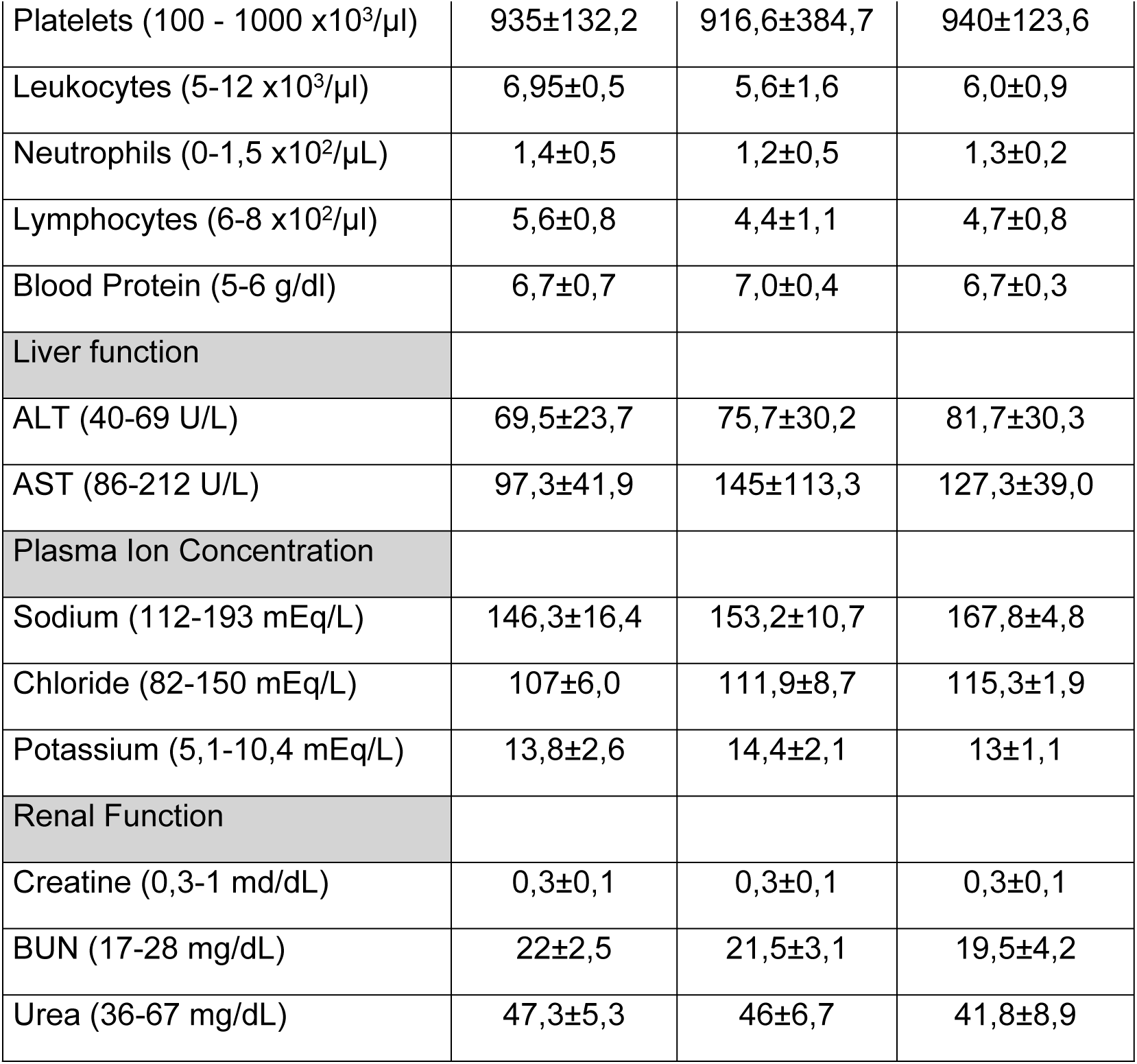
Blood count and chemistry values in mice across the three groups in the subchronic oral toxicity study. The control group received PBS, while the other groups received E-NP (empty nanoparticles) and OPN (oral nanoformulation with OBE100). Data are shown as means ± SD for each group. No statistically significant differences were observed in the measured parameters between the experimental groups. Reference values are from references 32 and 36.

An ophthalmological assessment of the anterior chamber was performed at the beginning and end of the study in mice from each treatment group to evaluate ocular toxicity related to 90-day administration. An examination of responses to stimuli, including proprioception, threat reflexes, and ocular reflexes (miosis/mydriasis), was also conducted. No signs of neurological or ocular toxicity were observed during the study.

On Day 14 post-treatment (PTD14), euthanasia was performed, and a necropsy was conducted. No differences in organ weights were observed, and macroscopically, no lesions were visible in any organ. Histopathological analyses were carried out on the liver, kidney, spleen, glandular and non-glandular stomach, pancreas, small intestine, and large intestine. No representative lesions were identified among the groups or associated with toxicity from the active ingredient or the vehicle (Fig. 4C).

### Combined study on chronic toxicity and carcinogenicity

One hundred twenty Swiss mice (CFW) were randomly assigned to three experimental groups, with twenty females and twenty males in each group for OPN, E-NP, and PBS controls: The OPN group received 300 μl (average 550 mg/kg) of the OBE100 nanoformulation; the E-NP group received 300 μl (average 460 mg/kg) of empty NPs; and the control group received 300 μl of PBS. The amount of oral nanoformulation and empty nanoparticles was halved compared to other treatments to facilitate administration, and the treatment period was extended to 10 months. Daily clinical monitoring showed no significant differences among the three groups. Body weight increased over the 10 months, with no differences among experimental groups or sex (Fig. 5A, B).

**Fig 5.**
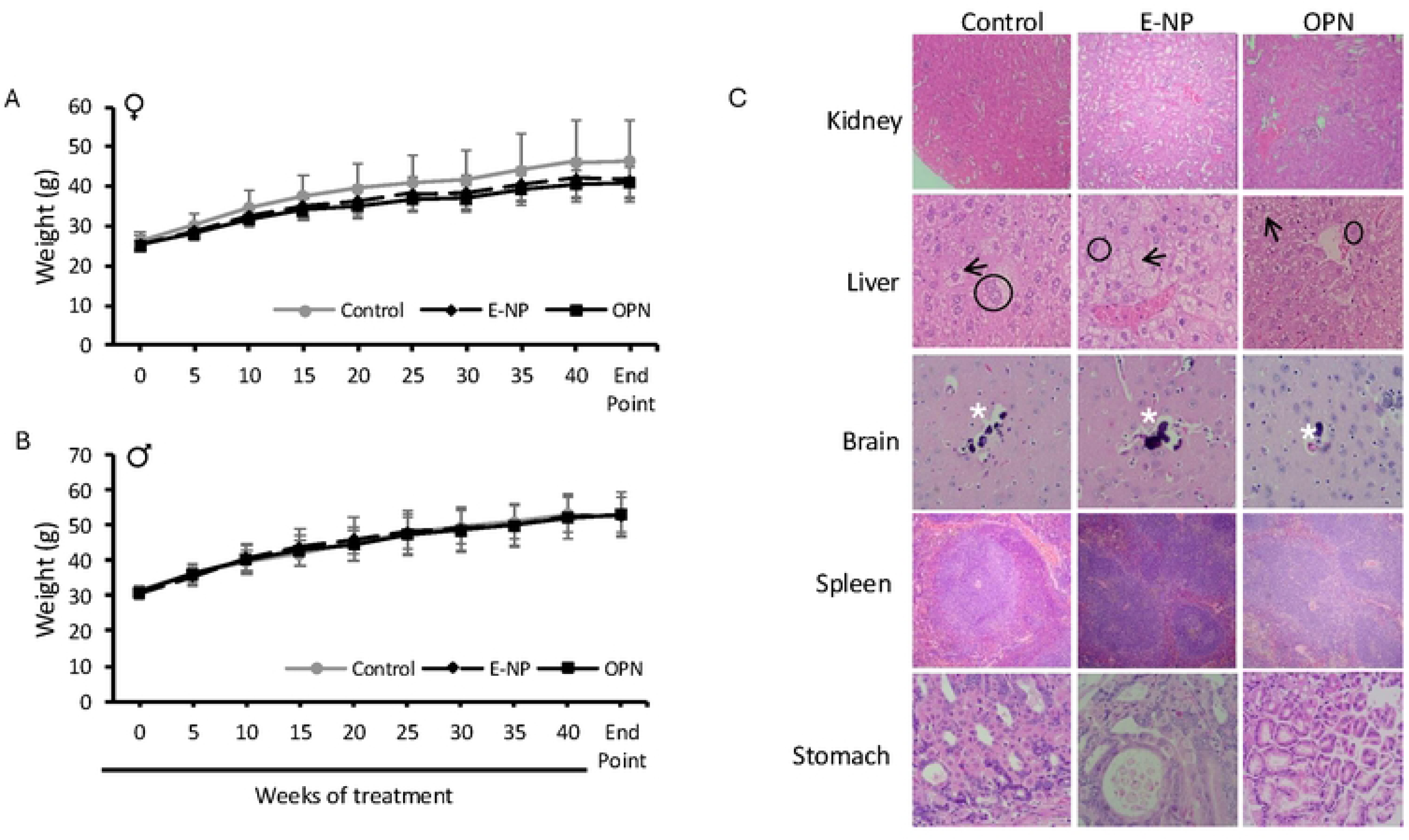
Body weight changes in the combined chronic toxicity and carcinogenicity study. A. Females, B. Males. C) Photomicrographs of the kidneys, liver, brain, spleen, and stomach from mice in three groups of the combined chronic toxicity and carcinogenicity study. The images are at 100X and 400X magnification and stained with hematoxylin and eosin (H&E). The control group received PBS; E-NP denotes empty nanoparticles; OPN indicates the oral nanoformulation with OBE100. **Liver**: Black arrow points to fatty degeneration; The black circle highlights binucleation. **Brain:** White asterisk marks mineral deposits.

Blood counts taken at the end of the study showed significant leukopenia, mainly lymphopenia, in all three trial groups. Clinical-pathological analyses of liver function revealed ALT levels well above the normal range for the control and E-NP groups (Table 3), with slightly elevated levels in the OPN-treated group. AST levels remained within the normal range. There is no pattern indicating liver toxicity caused by OPN. Overall, there is considerable variability across groups, as evidenced by large standard deviations, driven mainly by extreme values in a few animals. Alkaline phosphatase (ALT) levels are slightly elevated, while creatinine remains within the normal range (Table 3).

**Table 3.**
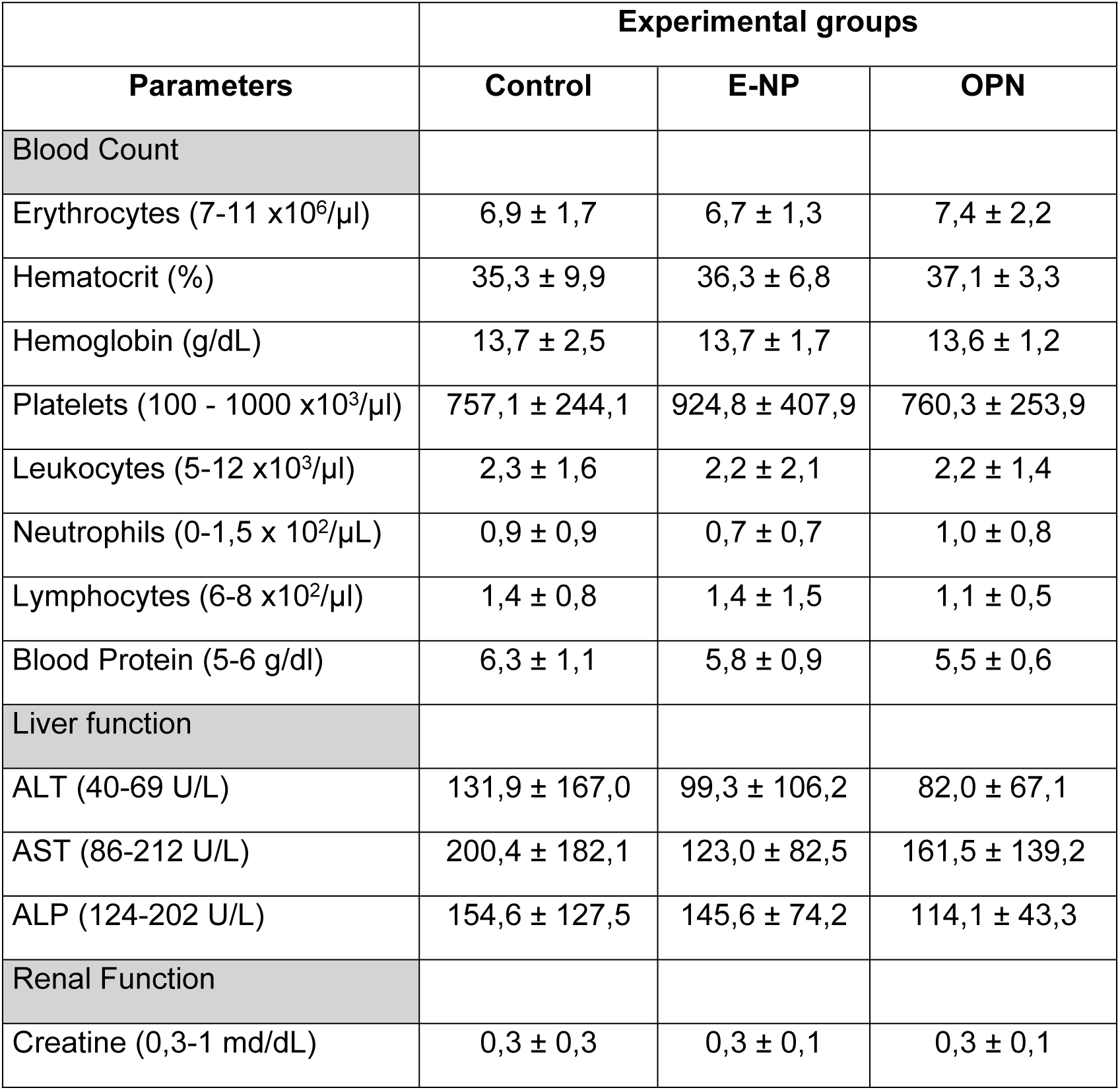
Blood count and chemistry values in mice across all three groups in a combined chronic toxicity and carcinogenicity study. The control group received PBS, while the other groups received E-NP (empty nanoparticles) or OPN (oral nanoformulation with OBE100). Data are presented as means ± SD for each group. No statistically significant differences were observed in the measured parameters between the experimental groups. Reference values are from references 32 and 36.

| Parameters | Control | E-NP | OPN |
| --- | --- | --- | --- |
| Blood Count |  |  |  |
| Erythrocytes (7-11 x10<sup>6</sup>/μl) | 6,9 ± 1,7 | 6,7 ± 1,3 | 7,4 ± 2,2 |
| Hematocrit (%) | 35,3 ± 9,9 | 36,3 ± 6,8 | 37,1 ± 3,3 |
| Hemoglobin (g/dL) | 13,7 ± 2,5 | 13,7 ± 1,7 | 13,6 ± 1,2 |
| Platelets (100 - 1000 x10<sup>3</sup>/μl) | 757,1 ± 244,1 | 924,8 ± 407,9 | 760,3 ± 253,9 |
| Leukocytes (5-12 x10<sup>3</sup>/μl) | 2,3 ± 1,6 | 2,2 ± 2,1 | 2,2 ± 1,4 |
| Neutrophils (0-1,5 x 10<sup>2</sup>/μL) | 0,9 ± 0,9 | 0,7 ± 0,7 | 1,0 ± 0,8 |
| Lymphocytes (6-8 x10<sup>2</sup>/μl) | 1,4 ± 0,8 | 1,4 ± 1,5 | 1,1 ± 0,5 |
| Blood Protein (5-6 g/dl) | 6,3 ± 1,1 | 5,8 ± 0,9 | 5,5 ± 0,6 |
| Liver function |  |  |  |
| ALT (40-69 U/L) | 131,9 ± 167,0 | 99,3 ± 106,2 | 82,0 ± 67,1 |
| AST (86-212 U/L) | 200,4 ± 182,1 | 123,0 ± 82,5 | 161,5 ± 139,2 |
| ALP (124-202 U/L) | 154,6 ± 127,5 | 145,6 ± 74,2 | 114,1 ± 43,3 |
| Renal Function |  |  |  |
| Creatine (0,3-1 md/dL) | 0,3 ± 0,3 | 0,3 ± 0,1 | 0,3 ± 0,1 |

For histopathological analysis, 12 of the 24 sampled mouse organs were excluded from the statistical analysis because they showed no lesions in any category. These were: cerebellum, adrenal gland, heart, eyes, ovaries, prostate, epididymis, testis, seminal vesicle, uterus, lymph node, and skin.

The organs showing characteristic histopathological lesions were as follows: 1. Spleen: atypical hyperplasia, more common in males and females, and lymphoid hyperplasia; 2. Brain: mineral deposits; 3. Esophagus: mucosal hyperplasia; 4. Glandular stomach: glandular dilation and hyperplasia, mucosal hyperplasia, increased populations of lymphocytes and plasma cells; 5. Non-glandular stomach: mild mucosal hyperplasia; 6. Salivary glands: mild chronic inflammation; liver: fatty changes, hydropic alterations, karyomegaly, binucleation, hypertrophy, and mild inflammatory foci; 7. Small intestine: mucosal hyperplasia, crypt hyperplasia, and GALT hyperplasia; 8. Kidneys: membranoproliferative glomerulonephritis and chronic interstitial inflammation (Fig. 5C). A semi-quantitative assessment of organ lesions was performed, and data on the most representative lesions are presented in Table 4.

**Table 4.** A semi-quantitative method was used to evaluate lesion severity in the histopathological analysis of organs from mice across all three groups in a combined chronic toxicity and carcinogenicity study. The control group received PBS, while the other groups received E-NP (empty nanoparticles) or OPN (oral nanoformulation with OBE100). Data are shown as means ± SD for each group. No statistically significant differences were observed in the measured parameters between the experimental groups.

| ORGANS/<br>ALTERATIONS | EXPERIMENTAL GROUPS |  |  |  |  |  |
| --- | --- | --- | --- | --- | --- | --- |
|  | Control |  | E-NP |  | OPN |  |
|  | F | M | F | M | F | M |
| **LIVER** |  |  |  |  |  |  |
| Binucleation | 1,5 ± 1,4 | 2,3 ± 0,6 | 1,5 ± 1,2 | 2,7 ± 0,8 | 1,9 ± 0,8 | 2,6 ± 0,8 |
| Fatty changes | 0,4 ± 1,0 | 0,9 ± 1,2 | 0,7 ± 1,1 | 0,5 ± 1,2 | 0,3 ± 0,8 | 1,0 ± 1,4 |
| Hypertrophy | 1,4 ± 1,9 | 1,8 ± 1,2 | 1,2 ± 1,3 | 2,6 ± 1,4 | 0,9 ± 1,0 | 2,4 ± 1,4 |
| Karyomegaly | 1,8 ± 1,3 | 3,1 ± 1,3 | 1,9 ± 1,1 | 2,9 ± 1,5 | 1,4 ± 0,8 | 2,6 ± 1,3 |
| Inflammatory Foci | 0,5 ± 0,7 | 0,2 ± 0,4 | 3,0 ± 1,4 | 1,0 ± 1,1 | 0,3 ± 0,6 | 0,7 ± 1,2 |
| **KIDNEY** |  |  |  |  |  |  |
| MG | 1,0 ± 1,1 | 1,2 ± 1,0 | 1,3 ± 1,4 | 1,3 ± 1,1 | 1,9 ± 1,1 | 0,6 ± 1,0 |
| Increased lymphocytes. | 1,5 ± 1,1 | 0,9 ± 1,0 | 1,1 ± 0,6 | 1,6 ± 1,2 | 1,3 ± 1,0 | 1,1 ± 1,1 |
| Increased Plasmacytes | 1,2 ± 0,8 | 0,9 ± 0,9 | 0,9 ± 0,4 | 1,4 ± 1,2 | 1,1 ± 0,7 | 0,9 ± 0,8 |
| Increased Macrophage | 1,0 ± 0,8 | 0,4 ± 0,5 | 0,7 ± 0,5 | 0,6 ± 0,7 | 1,1 ± 0,7 | 0,7 ± 0,6 |
| **SMALL INTESTINE** |  |  |  |  |  |  |
| Crypt hyperplasia | 0,8 ± 1,0 | 0,0 ± 0,0 | 0,5 ± 0,9 | 0,6 ± 1,1 | 0,3 ± 1,0 | 0,2 ± 0,8 |
| Mucosal hyperplasia | 0,5 ± 0,9 | 0,0 ± 0,0 | 0,7 ± 1,3 | 0,4 ± 1,0 | 0,2 ± 0,8 | 0,3 ± 0,8 |
| Villous atrophy | 1,2 ± 1,1 | 0,3 ± 0,7 | 0,6 ± 1,0 | 0,5 ± 1,0 | 0,3 ± 0,7 | 0,5 ± 1,0 |
| GALT hyperplasia | 1,3 ± 1,3 | 0,8 ± 0,8 | 1,4 ± 1,2 | 0,9 ± 1,4 | 1,1 ± 1,5 | 0,5 ± 0,8 |
| **GLANDULAR STOMACH** |  |  |  |  |  |  |
| Glandular dilatation | 1,4 ± 1,2 | 2,2 ± 1,4 | 1,4 ± 1,0 | 1,8 ± 1,5 | 1,5 ± 0,8 | 1,7 ± 1,2 |
| Glandular Hyperplasia | 0,4 ± 0,9 | 0,8 ± 1,1 | 0,4 ± 1,0 | 1,1 ± 1,4 | 0,1 ± 0,3 | 1,3 ± 1,2 |
| Mucosal Hyperplasia | 0,3 ± 0,7 | 1,2 ± 1,3 | 0,3 ± 0,5 | 0,3 ± 0,7 | 0,7 ± 1,2 | 0,9 ± 1,1 |
| Increased lymphocytes. | 0,3 ± 0,6 | 0,2 ± 0,6 | 0,1 ± 0,3 | 0,2 ± 0,5 | 0,2 ± 0,4 | 0,2 ± 0,6 |

MG: membranoproliferative glomerulonephritis
GALT: Gut-associated lymphoid tissues

In relevant organs and lesions, no statistically significant differences were observed between OPN, E-NP, and controls. Additionally, there were no consistent increases in severity (grade) or positivity (% > 0) attributed to OPN treatment after adjusting for multiple comparisons (Holm).

## Discussion

PLGA is a co-polymer known for its excellent biocompatibility, biodegradability, inherent non-toxicity, and superior encapsulation abilities [20]. It is used to produce nanoparticles that are widely used in preclinical research, addressing key limitations such as poor aqueous solubility, precipitation, and rapid systemic metabolism, which can decrease the therapeutic effectiveness of many natural extracts [21], including OBE100. In diabetes treatment, PLGA is often used for oral insulin delivery, providing protection from enzymatic degradation in the gastrointestinal tract and allowing controlled, sustained release [22]. Core-shell insulin/CS-PLGA nanoparticles showed relatively long-term hypoglycemic effects in diabetic rat models after oral administration [23]. Additionally, PLGA nanoformulations are crucial for encapsulating natural extracts like curcumin [24], Ƴ-Oryzanol [25], and resveratrol [26], as well as polyphenols and other compounds [27]. These studies confirm PLGA as a biocompatible platform that enhances the pharmacological potential of natural compounds, thereby improving therapeutic outcomes in preclinical trials, as demonstrated by the platform used here to encapsulate the OBE100 extract.

For the preclinical trial of the PLGA nanoformulation, the standardized scale-up method described here for producing reproducible OPN from 8 mL to 500 mL using the emulsion/solvent-evaporation technique maintained the critical quality attributes (size, polydispersity, and ζ-potential) despite roughly a 60-fold increase in volume. This result shows that careful control of processing parameters can reduce the inherent heterogeneities in mass and energy transfer typically encountered during scaling [28], as observed in previous studies on the scaling up of encapsulated natural compounds such as curcumin using PLGA [29]. While earlier research often recommends switching to advanced homogenization techniques such as high-pressure homogenization or inline sonication for large-scale PLGA production to ensure consistency and high throughput [28], our data and scale-up method demonstrate that standard homogenization—by controlling sonication time, pulsing, and power—can produce successful results while maintaining the essential process parameters. This intermediate scale-up, even with conventional sonication, suggests that the specific formulation and the method described for scale-up may provide a degree of process robustness, but moving to pilot scale (> 1 L) will likely require adopting continuous-flow systems to achieve industrial scalability and batch-to-batch consistency [30].

In acute and subacute toxicity tests, mice showed no significant clinical changes or macroscopic findings, and histopathological analyses of organs indicated that the developed nanoformulation is safe, with no toxicity observed under the study conditions.

Blood chemistry analyses in mice during the subchronic toxicity study showed that only AST was slightly elevated in the OPN-treated group compared to the control group; however, this difference was not significant for organ function. Although liver disease is usually suspected initially based on elevated liver enzyme activity in screening profiles, ALT and AST are sensitive biomarkers of hepatocellular function, and their results should be correlated with macroscopic findings, organ weight, and histopathological results. It is also important to assess the level of enzyme activity elevation relative to the reference range: mild (<3-fold), moderate (3-9-fold), and marked (>10-fold) [31]. In our study, ALT levels in the control and study groups and AST levels in the OPN group were mildly elevated, with no significant tissue changes observed. Furthermore, the comparison between the experimental groups revealed no statistically significant differences. When we repeated the analysis in the subchronic study, transaminase levels remained within the normal range. Given that this study involved a larger number of animals over a longer period, it supports the compound’s safety at the hepatic level.

The discrepancy between the studies may be due to differences in the number of mice per group, leading to a higher standard deviation in the subacute study.

Subchronic oral toxicity studies confirmed that OPN does not cause death or signs of toxicity in mice, indicating its potential for safe use. We observed that the control group mice gained more weight than the other two groups (Fig. 4A, B). This is likely due to the smaller orogastric tube used in the control group, which reduced the stress the mice experienced during treatment.

Although daily clinical monitoring showed no significant changes across the three experimental groups, the combined chronic toxicity and carcinogenicity study revealed marked leukopenia, particularly lymphopenia, at the end of the study in all three test groups (Tables 3 and 4). Studies on adult Swiss/ICR mice, an outbred strain, some of which are stratified by age, generally indicate that, with age, the total white blood cell count decreases, the percentage of lymphocytes declines, and the percentage of neutrophils increases [32]. Research has also documented a gradual reduction in lymphocyte and monocyte counts during periods of stress [33]. Since the PBS-treated control group also exhibited levels below the normal range, this parameter is not attributed to the active ingredient or the vehicle. No other changes were observed in the red line. Additionally, pulmonary adenomas were seen in all three experimental groups, but no neoplastic proliferative disorders related to the treatment or vehicle were observed. This type of benign tumor develops spontaneously in Swiss mice aged 6 months or older [34, 35].

Analysis of liver function showed elevated ALT levels in the control and E-NP groups, above the normal range (Table 3); however, the OPN-treated group had the lowest levels, closest to normal. AST and ALP remained within normal limits. Overall, there is significant variability among individuals, as indicated by the high standard deviations. Previous studies have shown that AST varies by strain and sex and is not highly liver-specific in rodents [34].

In relevant organs and lesions, no statistically significant differences were observed among the experimental groups, nor were there consistent increases in severity (grade) or positivity (%>0) attributed to OBE-100 after adjustment for multiple comparisons (Holm). The observed patterns align with background biological variability, reactive or regenerative changes, or effects unrelated to the active ingredient. Based on these data, no treatment-related histopathological signals that would prevent progression to clinical studies are identified, supporting continued targeted monitoring.

Building on this information, adaptive changes such as hyperplasia of the esophageal mucosa and non-glandular stomach may have resulted from repeated trauma caused by the orogastric tube used in daily treatments. Lymphoplasmacytic infiltrates in the digestive tract, observed in all three groups, are consistent with responses to luminal or microbiota antigenic stimuli.

In the liver, polyploidy, binucleation, and hypertrophy—changes not linked to any specific group—increase with age in rodents and are associated with genetic background lesions. Hydropic change correlates with osmotic-ischemic alterations or postmortem changes. Fatty change was also not significantly more common in the E-NP and OPN groups and may be related to other metabolic or nutritional factors. In the kidney, membranoproliferative glomerulonephritis is also a mild genetic background lesion and is not caused by treatment.

In toxicological studies involving rodents, the term “atypical hyperplasia” almost always refers to reactive lymphoid hyperplasia with less typical features, such as immense germinal centers, prominent marginal zones, or slight expansion of the white pulp. However, it maintains the typical architecture of the spleen, including the white and red pulp. This preservation of structure is a key feature that distinguishes lymphoid hyperplasia; INHAND/NTP emphasizes that, in lymphoid hyperplasia, splenic organization remains intact, whereas in lymphoma it becomes distorted [35]. Splenic lymphoid hyperplasia in mice is common and usually mild, typically associated with antigenic stimulation and often characterized by increased cellularity in follicles, germinal centers, and the marginal zones.

Since this lesion was consistent across all three groups and showed no sex difference, the diagnosis is reactive lymphoid hyperplasia of genetic origin rather than a compound-related event.

There is no specific clustering of tumors in OPN: pulmonary adenomas, common background lesions in aging mice, appear in both the control and E-NP groups; adenocarcinomas are observed in E-NP; lymphomas occur in both OPN and E-NP; and gastric carcinoma is unique, with no consistent pattern. This distribution aligns with the expected background in Swiss mice and does not indicate an oncogenic effect from OBE100.

## Conclusion

Preclinical in vivo studies conducted in accordance with OECD guidelines, including assessments of acute, subacute, subchronic, and chronic/carcinogenicity, showed no treatment- or vehicle-related toxicity, supporting its safety in mice. This conclusion is based on clinical and paraclinical data. No histopathological lesions, such as degeneration or cell death, were observed in the liver, kidney, or gastrointestinal tract. Similarly, the respiratory, nervous, and reproductive systems were unaffected by the treatment. Clinical pathology studies, including blood chemistry and complete blood counts, further reinforce these findings.

## Author contributions

Writing - original draft, investigation, formal analysis, methodology, visualization (Natalia Arbelaez, Elkin Escobar-Chaves). Investigation, formal analysis, validation (Sergio Acin, Andrea Correa, Adriana Restrepo). Conceptualization, formal analyses, Writing - Review & Editing (Jahir Orozco). Conceptualization, formal analyses, writing - Review & Editing, visualization, supervision, project administration, and funding acquisition (Norman Balcazar).

## Ethics approval

The Institutional Animal Care and Use Committee of the University of Antioquia approved all animal experimental procedures.

